# Biochemical characterization of a human septin octamer

**DOI:** 10.1101/2021.09.06.459054

**Authors:** Martin Fischer, Dominik Frank, Reinhild Rösler, Nils Johnsson, Thomas Gronemeyer

## Abstract

Septins are part of the cytoskeleton and polymerize into non-polar filaments of heteromeric hexamers or octamers. They belong to the class of P-loop GTPases but the roles of GTP binding and hydrolysis on filament formation and dynamics are not well understood.

The basic human septin building block is the septin rod, a hetero-octamer composed of SEPT2, SEPT6, SEPT7, and SEPT9 with a stoichiometry of 2:2:2:2 (2-7-6-9-9-6-7-2).

Septin rods polymerize by end-to-end and lateral joining into linear filaments and higher ordered structures such as rings, sheets, and gauzes.

We purified a recombinant human septin octamer from *E. coli* for *in vitro* experimentation that is able to polymerize into filaments. We could show that the C-terminal region of the central SEPT9 subunit contributes to filament formation and that the human septin rod decreases the rate of *in vitro* actin polymerization.

We provide further first kinetic data on the nucleotide uptake- and exchange properties of human hexameric and octameric septin rods. We could show that nucleotide uptake prior to hydrolysis is a dynamic process and that a bound nucleotide is exchangeable. However, the hydrolyzed γ-phosphate is not released from the native protein complex. We consequently propose that GTP hydrolysis in human septins does not follow the typical mechanism known from other small GTPases.

## 2 Introduction

Septins were discovered in yeast in 1974 and are part of the cytoskeleton (Mostowy and Cossart 2012). They form non-polar filaments that unlike actin filaments and microtubules do not serve as railroad tracks for intracellular transport. Being rather neglected for two decades after their discovery, research on the septins gained a growing attention over the past few years (Menon, M. B., Gaestel 2017). Initially labeled as passive scaffold proteins, septins were later discovered to actively participate in many dynamic intracellular processes. Septins regulate the organization of the cytoskeleton, vesicle transport and fusion, chromosome alignment and segregation, and cytokinesis (Surka, Tsang, and Trimble 2002; Estey et al. 2010; Bowen et al. 2011; Fuechtbauer et al. 2011; Tokhtaeva et al. 2015). The septin subunit SEPT9 was reported to bundle actin and microtubuli (Mavrakis et al. 2014; Dolat et al. 2014; Bai et al. 2013; Smith, Dolat, Angelis, Forgacs, Spiliotis, Galkin, et al. 2015).

The mammalian genome encodes thirteen different septins (SEPT1-SEPT12, SEPT14) (Hilary Russell and Hall 2011), of which SEPT2, SEPT7 and SEPT9 are nearly ubiquitously expressed, while SEPT1, SEPT3, SEPT12, and SEPT14 are tissue-specific. All septin subunits can be sorted into four subgroups based on sequence homology, namely the SEPT2, SEPT3, SEPT6 and SEPT7 subgroup (Neubauer and Zieger 2017).

The basic septin building block in mammalian cells is a hetero-octamer composed of the SEPT2, SEPT6, SEPT7, and SEPT9 at a stoichiometry of 2:2:2:2 (2-7-6-9-9-6-7-2) (Soroor et al. 2020). This building block is capable of polymerizing by end-to-end and lateral joining into higher ordered structures such as rings, filaments, and gauzes. Assembly into these higher ordered structures is maintained by alternating interactions between the G domains (G interface) or between the N- and C-termini (NC interface), respectively (Bertin et al. 2008).

All septins share a central GTP binding domain (short G domain) that is flanked by variable C- and N-terminal extensions (Sirajuddin et al. 2007) and contains a loop with two distinct beta sheets, the “septin unique element”. The G domain is conserved among different species and is supposed to be loaded either with GTP or GDP. The available crystal structures of human and yeast septins (Zent, Vetter, and Wittinghofer 2011; Sirajuddin et al. 2007, 2009; Brausemann et al. 2016; Castro et al. 2020; Fonseca Valadares et al. 2017; Rosa et al. 2020) confirm that the G domain contains all structural features of small GTPases like the P-loop and the distinct Switch1 and Switch2 loops including the invariant DXXG motif. GTP binding and hydrolysis was confirmed for human septins. Standard assays aiming at detecting GTP hydrolysis in small GTPases employ usually loading of the GTPase with a GTP labeled at its γ-phosphate and subsequent detection of the released γ-phosphate (Frech et al. 1990) or detection of GTP decrease and GDP increase upon hydrolysis by HPLC (Tucker et al. 1986). However, detection of GTPase hydrolysis products in human septin subunits was so far only achieved after denaturation of the protein (Castro et al. 2020; Sirajuddin et al. 2009). GTP uptake and hydrolysis of entire rods was not yet examined.

Only one of the available septin crystal structures, the human SEPT2/6/7 complex (Sirajuddin et al. 2007; Mendonça et al. 2021), shows a rod-like structure. All other structures represent septin monomers or dimers. An octameric rod for *in vitro* experimentation was only presented recently (Iv et al. 2021) and in this work. We present here a first biochemical characterization of the nucleotide binding- and hydrolysis properties of purified human octameric septin rods as well as a kinetic assay that allows to measure the influence of septin complexes on actin filament assembly in real time.

## 3 Materials and Methods

### Molecular cloning, protein purification- and analysis

The ORFs of the isoforms i1 of SEPT2 and SEPT9 (or SEPT9_1-568_) as well as SEPT7 and SEPT6 were cloned into two compatible bicistronic plasmids. SEPT6 was constructed with a N-terminal 6His-tag for IMAC purification while the other subunits were untagged.

Protein expression of hexameric and octameric rods was performed in the *E. coli* strain BL21DE3 in SB medium at 18 °C. Protein purification from the crude extract was performed by immobilized metal affinity chromatography (IMAC) and subsequent size exclusion chromatography.

Integrity of the purified complex was determined by density gradient centrifugation and MS analysis. Detailed protocols on the cloning-expression and purification procedures as well as on the analytical assays can be found in the supporting methods.

### Filament formation assays

Septin filament formation was performed as described elsewhere for yeast septins (Renz, Johnsson, and Gronemeyer 2013). Briefly, septin preparations were adjusted to 2 μM and cleared from aggregates by centrifugation. Filament formation was induced by dialysis of hexameric and octameric rods into buffers with various NaCl concentrations, followed by incubation at 4 °C overnight and subsequent centrifugation at 100.000 xg.

### Nucleotide uptake- and hydrolysis assays

Uptake of [γ^32^P]-GTP to hexameric and octameric rods and isolated SEPT9 was performed as described elsewhere for yeast septins (Baur et al. 2018). Briefly, septin preparations were incubated with [γ^32^P]-GTP in exchange buffer containing 5 mM EDTA. For hydrolysis, the exchange reaction was transferred into hydrolysis buffer containing MgCl_2_. Radioactivity retained in the native septin proteins during uptake or during hydrolysis was detected with a filter binding assay.

Further details of the procedure and the subsequent data evaluation are outlined in the supporting methods.

### Actin polymerization assays

Pyrene-labeled G-actin (shortly pyrene actin) was prepared from rabbit muscle acetone powder using an already described protocol (Cooper, Walker, and Pollard 1983) with some modifications. Pyrene actin fluorescence increases upon actin polymerization and can be recorded in actin polymerization buffer at an excitation of 365 nm and emission of 407 nm.

Details on the pyrene actin preparation and the assay protocol are outlined in the supporting methods.

## 4 Results

### Construction and purification of human hexameric- and octameric septin rods

We expressed human octameric and hexameric septin rods in *E.coli* from two compatible bicistronic plasmids. We followed the subunit combination on the plasmids chosen by a previous study (Sheffield et al. 2003): SEPT6 and SEPT7 were expressed from one plasmid and SEPT2 (and SEPT9 for octameric rods) from the second plasmid. In our setup only SEPT6 carried a hexa-histidine tag for purification by immobilized metal affinity chromatography (IMAC). The enriched fractions were subsequently subjected to size exclusion chromatography (SEC). SDS-PAGE documented the presence of shorter fragments of SEPT9 that could not be separated from the rods during the purification. (Figure1A). To avoid the generation of these fragments we truncated the SEPT9 subunit at Glycine 568 (SEPT9_G568_). Glycine 568 represents the terminal residue α6 helix of the G domain and is also the terminal residue of the isoform SEPT9i3. Octameric rods expressing SEPT9_G568_ showed less degradation products than the native octamer and the purified complex could be readily separated from other subcomplexes and contaminants by the preparative SEC (Figure 1B). The purified complex eluted in a shoulder of the main product peak (Suppl. Figure 1A).

**Figure 1.**
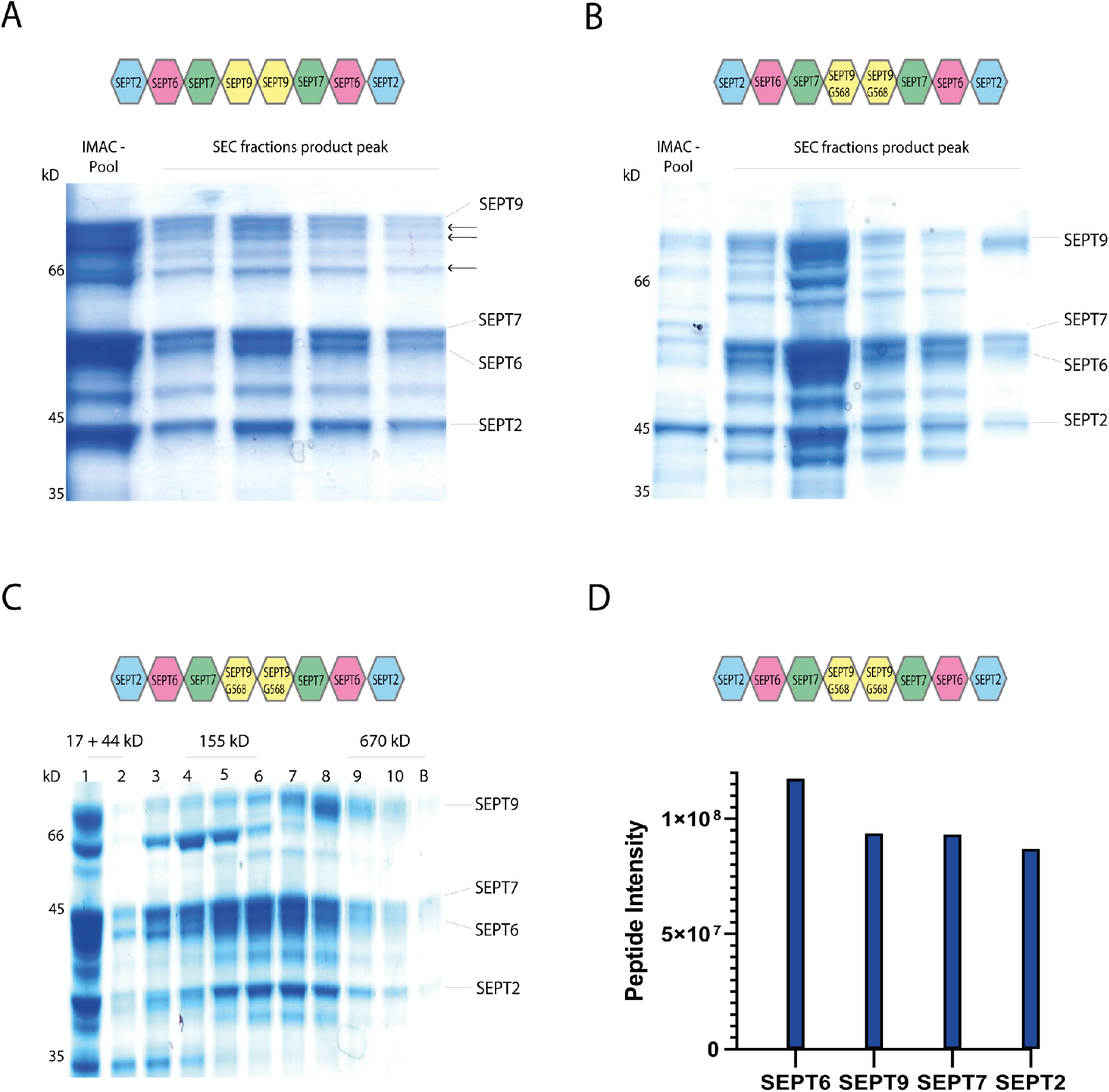
Purification of octameric septin rods. **A.** SDS-PAGE (Coomassie staining) of a representative purification of a septin octamer containing SEPT9_FL_. IMAC and SEC fractions are indicated. SEPT9 degradation products are marked with arrows. **B.** SDS-PAGE (Coomassie staining) of a representative purification of a septin octamer containing SEPT9_G568_. IMAC and SEC fractions are indicated. A chromatogram plot of the SEC is shown in Suppl. Fig. 1A. **C.** SDS-PAGE (Coomassie staining) of the fractions from a 15% - 30% Glycerol gradient separation of the SEPT9_G568_ octamer. The kD of the corresponding molecular weight marker is indicated. The separation of the marker proteins is shown in Suppl. Fig. 1B. **D.** Peptide abundancies of the SEPT9_G568_ octamer determined by MS analysis. The intensity score reflects the accumulated intensities of all peptides detected for the respected subunit.

The enriched septin preparation from IMAC was analysed on a Glycerol gradient centrifugation to confirm the integrity of the complex. Septin subunits, SEPT9 degradation products and contaminants migrated in the fractions corresponding to a lower molecular weight whereas the combination of all four subunits migrated just before the 670 kD marker protein became abundant, indicative of an octameric septin complex (Figure 1C, Suppl. Figure 1B).

MS of the purified complex revealed that all four subunits are present in nearly stoichiometric abundance (Figure 1D). Tagged subunits of complex preparations are usually more abundant than untagged ones which is also the case for SEPT6. We conclude that our septin preparation is indeed an octameric complex containing all four subunits.

Hexameric SEPT2/6/7 complexes were purified using the same protocol (Suppl. Fig 1C.).

### Filament forming properties

We asked next whether septin filament formation is altered by the absence of the SEP9 C-terminal extension. Septin preparations were cleared from aggregates by centrifugation, dialyzed into low salt buffer and subjected subsequently to ultracentrifugation. Surprisingly, we could detect filaments even at 300 mM NaCl for SEPT9_FL_ octamers (Figure 2A). Filaments of SEPT9_G568_ containing octamers and hexamers could be detected in the pellet at 50 mM NaCl whereas almost no filament formation occurred at 100 mM NaCl (Figure 2B, B). The removed C-terminal amino acids of SEPT9 follow the terminal α6 helix of the G domain which contributes to an interaction network with the neighboring subunit in the NC interface (Figure 2D). This interface might be stabilized by the terminal amino acids (see below).

**Figure 2.**
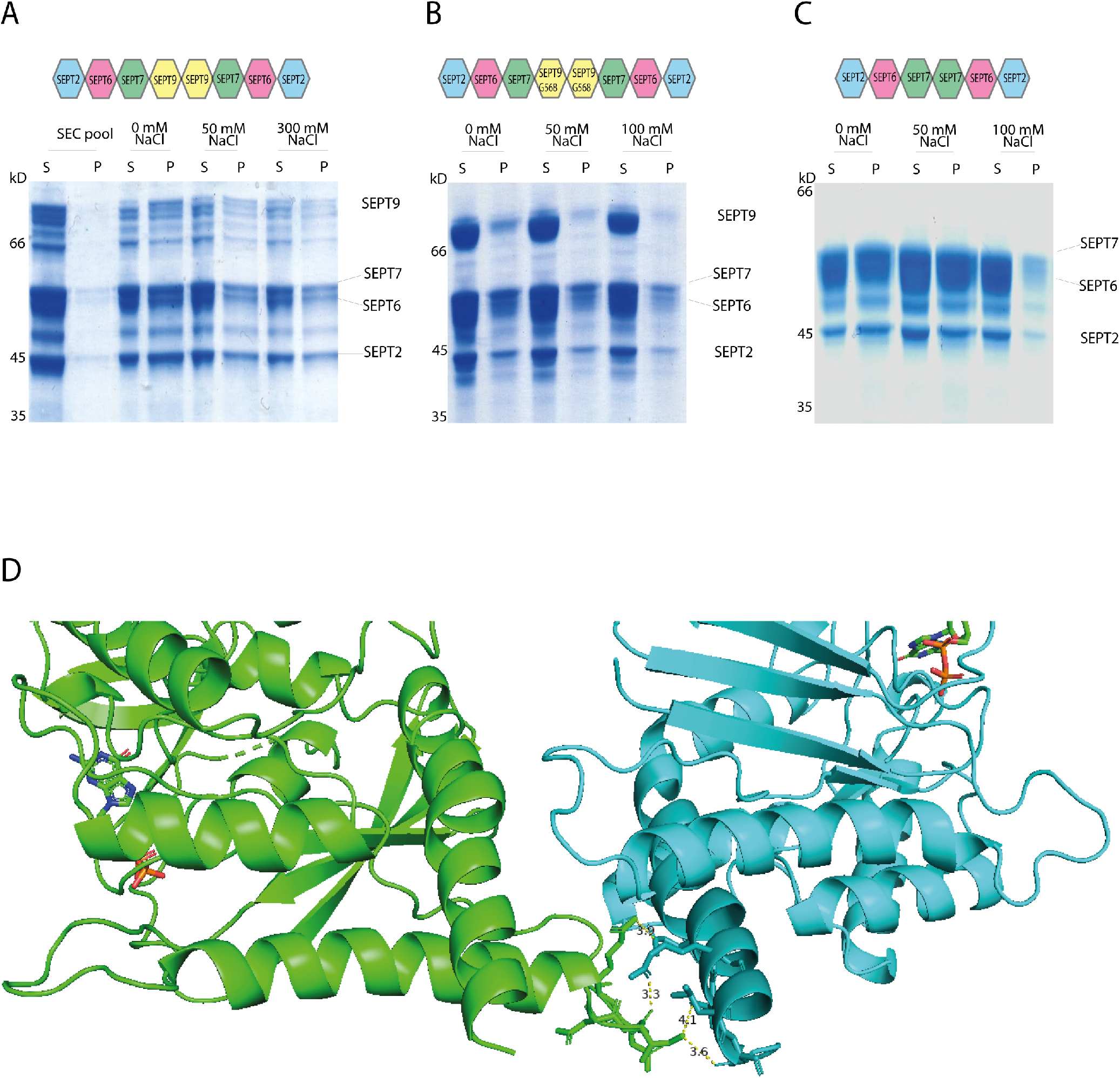
Filament formation properties of hexameric - and octameric septin rods. **A.** Filament formation of SEPT9_FL_ containing rods detected by a spin down assay. In SEC buffer containing 500 mM NaCl no filaments could be detected in the pellet (SEC pool). After dialysis and subsequent sedimentation, filaments could be detected in the pellet at all tested NaCl concentrations. S – supernatant, P – pellet. **B, C.** Filament formation of SEPT9_G568_ containing rods (B) and hexameric rods (C) detected by a spin down assay. After dialysis and subsequent sedimentation, filaments could be detected in the pellet at 0 mM and 50 mM NaCl whereas filament formation was almost abolished at 100 mM NaCl. **D.** Detail of the NC interface between two SEPT9 subunits (GDP-SEPT9; open NC-interface. PDB-ID 5CYO). Amino acids contributing to the interaction network between the α6 helix (dark-cyan) of one subunit (cyan) and the neighboring subunit (green) are highlighted.

### Effects of human septin rods on the polymerization properties of actin

The septin subunit SEPT9 was reported to bundle actin and microtubuli (Mavrakis et al. 2014; Dolat et al. 2014; Bai et al. 2013; Smith, Dolat, Angelis, Forgacs, Spiliotis, Galkin, et al. 2015).

To investigate the influence of septin rods on the formation of the actin cytosceleton *in vitro*, we performed actin polymerization assays with 2 μM pyrene labeled G-actin and recorded the increase of fluorescence over time. Without additives, a typical sigmoidal polymerization curve was obtained (Fig. 3A and B) (Cooper, Walker, and Pollard 1983) which can mathematically described by a growth curve including lag phase, exponential growth and plateau phase. From this curve, parameters such as the rate constant, the lag time or the doubling time can be calculated. All parameters calculated from the polymerization curves are summarized in Suppl. Table 1. Actin polymerization did not occur in G-buffer without salts (Suppl. Fig. 2A). Hexameric septins increased the lag time leading to a shift of the inflection point relative to the inflection point of the actin curve by 381 s (0.5 μM septin) and 1410 s (1 μM septin), respectively (Fig. 3A).

**Figure 3.**
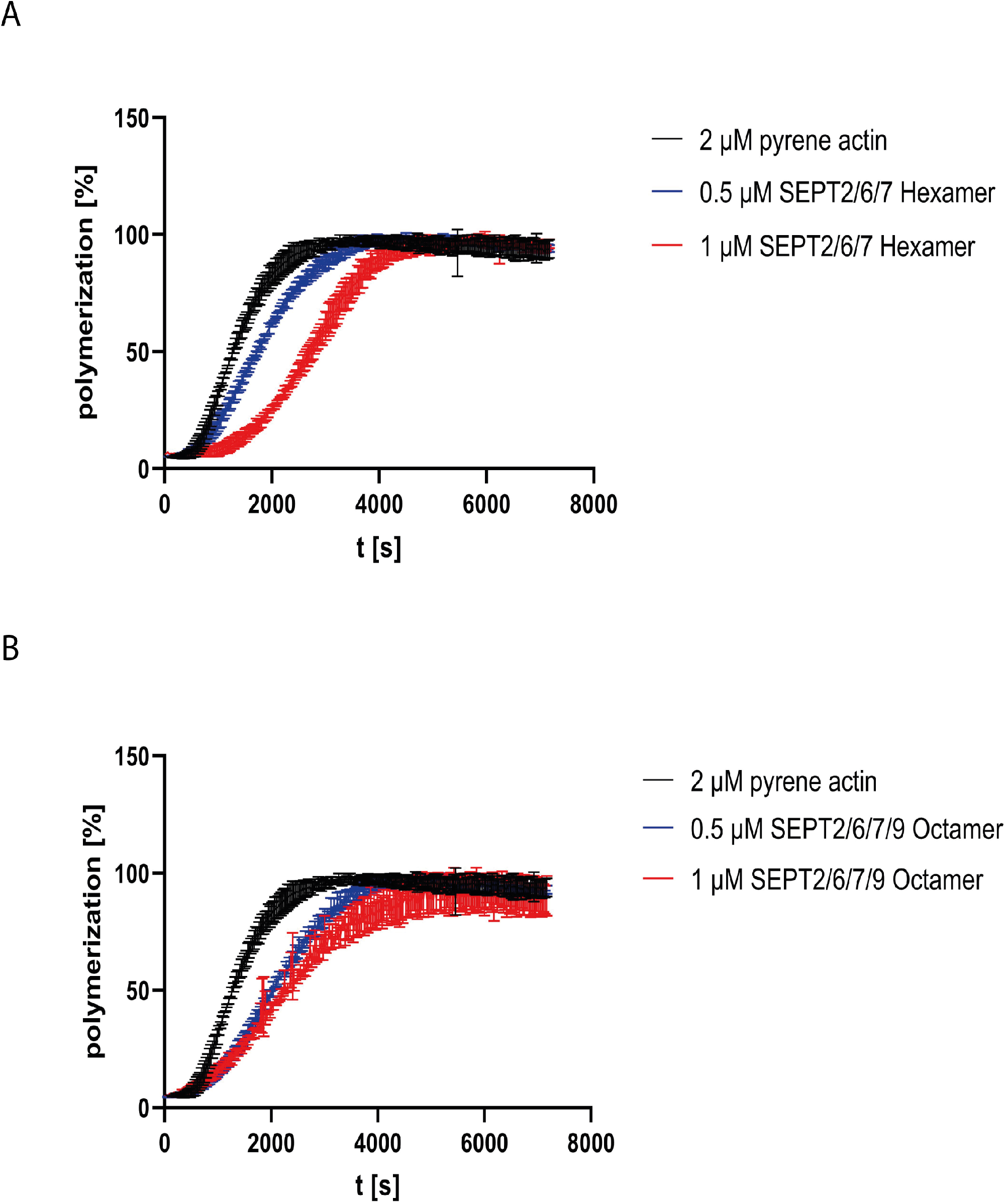
Influence of septin complexes on actin polymerization. Actin polymerization assay using pyrene actin with and without septin hexamers (A) or SEPT9_G568_ containing octamers (B). All assays were performed at least in triplicate.

For octamers, the shift was more pronounced by 637 s (0.5 μM septin) but did not change significantly at higher septin concentrations (724 s for 1 μM septin) (Fig. 3B). The actin monomer turnover in the elongation phase (represented by *τ*, the inverse value of the rate constant) was slightly decreased about 1.4 fold for hexameric rods and 0.5 μM octamer relative to actin alone and more pronounced by 2.1 fold for 1 μM octamer. Increased *τ* values are reflected by a shallower slope of the curve.

A spin down assay with septin octamers previously dialyzed in G buffer showed that septin filaments did not measurably bind G-actin (Suppl. Fig. 2B).

### Nucleotide uptake and hydrolysis properties of the human septins

We have previously measured the uptake of [γ^32^P] labelled GTP in the presence of EDTA by the hexameric- and octameric septin rods from yeast (Baur et al. 2018). We subjected our human septin rods to the same assay. All experiments were conducted with SEPT9_G568_ containing octamers.

[γ^32^P]-GTP uptake followed an exponential one phase association kinetics from which the rate constant and the reaction half time t_1/2_ can be calculated. The reaction half time t_1/2_ was 10 min for hexameric rods and 16 min for octameric rods, respectively (Fig. 4A). All t_1/2_ values of this and the subsequently performed experiments are summarized in Fig. 4C and the full evaluation of all performed measurements are shown in the supporting information (Suppl. Tables 2-4).

**Figure 4.**
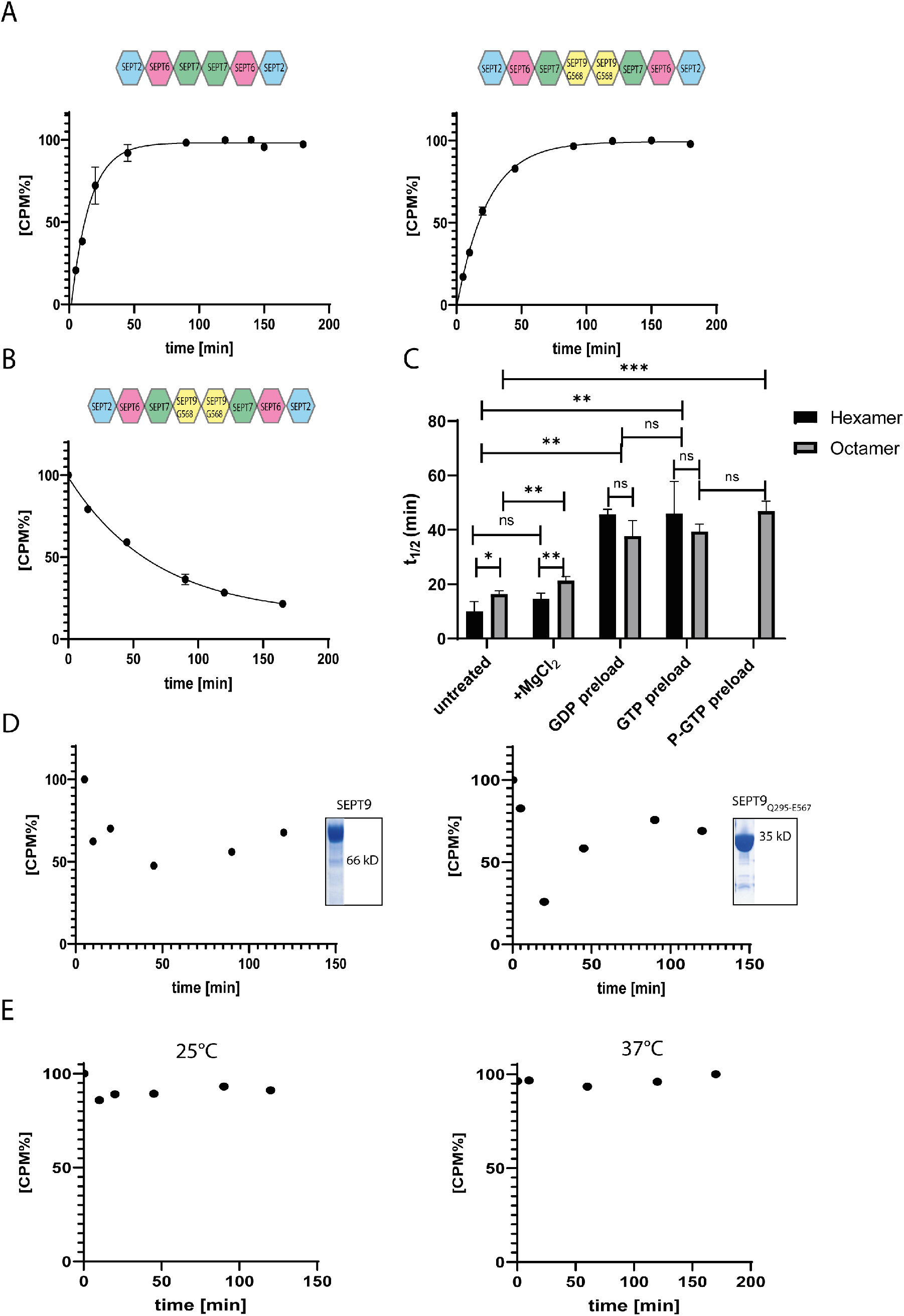
Nucleotide uptake- and hydrolysis properties of hexameric and SEPT9_G568_ octameric septin rods. **A.** Nucleotide exchange reaction of hexameric (left panel) and octameric septins (right panel). Purified complexes were incubated with [γ^32^P]-GTP and uptake was monitored at the indicated timepoints by a filter assay. [CPM%] values (normalized radioactive counts) are plotted vs. the reaction time. The connecting line represents the fitting curve of the exponential association. **B.** Nucleotide exchange reaction of octameric septins preloaded with [γ^32^P]-GTP. The preloaded complex was incubated with GTP and decrease of [γ^32^P]-GTP was monitored at the indicated timepoints by a filter assay as in A. The connecting line represents the fitting curve of the exponential dissociation. **C.** Compilation of the t_1/2_ values of all performed nucleotide exchange assays. The parameters of the statistical evaluation are provided in Supplementary Tables 2 and 3. Error bars represent the standard deviation. All assays (including those depicted in detail in A and B) were performed at least in triplicate. **D.** Representative [γ^32^P]-GTP uptake assay for isolated SEPT9_FL_ (left panel) and SEPT9_Q295-E567_ (right panel) subunits. The assay was performed as in A. The inlets show the Coomassie stained SDS-PAGE of both purifications. **E.** Representative GTP hydrolysis assay for octameric septins. After preloading with [γ^32^P]-GTP, the complex was subjected to hydrolysis conditions applicable for small GTPases. The reaction was performed at 25°C (left panel) and 37°C (right panel). Samples were taken at the indicated time points and the filter assay was performed as in A. GTP hydroloysis of hexameric rods is shown in Suppl. Fig. 4.

Adding MgCl_2_ to the exchange reaction after 20 min did not change the overall shape of the uptake kinetics but prolonged t_1/2_ slightly (and barely significantly) to 14.5 min for hexamers and 21 min for octamers (Suppl. Fig. 3A and Fig. 4C).

To evaluate whether the bound nucleotide remains in the septin rod or exchanges with free GTP, we pre-loaded septin preparations with non-radioactive GDP or GTP and removed excess nucleotide by micro-dialysis. Subsequent nucleotide exchange against [γ^32^P]-GTP showed a shift in reaction half times to 42 min for both GDP and GTP loaded hexameric rods and to 33 min for GDP loaded octameric rods and 38 min for GTP loaded octameric rods, respectively (Suppl. Fig. 3 B and C and Fig. 4C).

We also subjected the octameric rods to exchange of previously bound [γ^32^P]-GTP against non-radioactive nucleotide. After loading, excess nucleotide was removed via a desalting column and the uptake of non-radioactive GTP was monitored via the decrease of protein-bound radioactivity. The calculated reaction half time of 46.5 min was slightly (and not significantly) slower than the t_1/2_ for the inverse exchange reaction (GTP against [γ^32^P]-GTP) (Fig. 4B).

We conclude that human septin octamers exchange bound nucleotides quite readily.

Previous studies showed that isolated septin subunits may exhibit GTPase properties than the rods (Sheffield et al. 2003). We tested the isolated SEPT9_FL_ subunit and the SEPT9 G domain (SEPT9_Q295-E567_) for nucleotide uptake and found that both did not accept [γ^32^P]-GTP (Figure 4D).

To find out whether human septins follow the standard GTP hydrolysis mechanism than other small GTPases we measured the release or protein-bound radioactivity by [γ^32^P]-GTP preloaded septins under conditions used for other small GTPases (i.e. magnesium and an excess of non-radioactive GTP to avoid unwanted re-association of [γ^32^P]-GTP molecules) (Tucker et al. 1986; Schweins et al. 1995). In contrast to the classical small GTPases, neither hexameric nor octameric rods showed a release of hydrolyzed [γ^32^P] under these conditions, regardless of the applied reaction temperature (25 °C or 37 °C, respectively) (Fig. 4E and Suppl. Fig. 4).

This is more in line with our findings for yeast septins (Baur et al. 2018) rather than with the properties of small GTPases of the Ras family.

## 5 Discussion

### The C-terminus of SEPT9 contributes to filament formation

Human hexameric septin rods are available for in vitro experimentation and even crystallization since several years (Sheffield et al. 2003; Sirajuddin et al. 2007). However, purified octameric rods containing also SEPT9 were only very recently obtained (Iv et al. 2021). We introduce here an additional simple, two step purification protocol for human octameric rods harboring a hexa-histidine tag at only one of its subunits. Gycerol density gradient centrifugation of the complex after the first purification step separated contaminants and septin containing subcomplexes from the octamer, which migrated at the expected density in the gradient. After preparative SEC, we obtained the purified octamer including all four subunits in stoichiometric abundance.

The central subunit SEPT9 is prone to C-terminal degradation during expression and purification of the rod. Removal of the terminal 18 amino acids stabilized the protein and led to less degradation products. To our surprise we found that the SEPT9_FL_ containing rods formed filaments already at 300 mM NaCl whereas for hexamers and SEPT9_G568_ containing filament formation was abolished already at 100 mM NaCl. The first study of filament formation of SEPT2/6/7 hexamers raised the expectation of another – at that time largely unknown - septin subunit (SEPT9) with a stabilizing effect on the filaments (Sheffield et al. 2003).

The comparison of the crystal structures of SEPT9 in its GDP and GTPγS loaded state (PDB-IDs 5CYO and 5CYP) provides an explanation for our findings:

SEPT9 is the central subunit of the octameric rod. The α6 helix of the SEPT9 G domain is an essential component of the central SEP9-SEPT9 NC interface, contributing to an interaction network with the α2 helix of the neighboring subunit (Castro et al. 2020) (Fig. 2D). The 18 amino acids following the terminal G568 residue of the α6 helix are not covered by crystal structure but it can be anticipated that this flexible tail region stabilizes the position of the α6 helix in the interface. The tail contains five glutamic acid residues and a single lysine that may interact with the negatively charged sidechains of the neighboring tail. Without the presumably interacting tails, the helix might become more flexible, thereby disturbing the interaction network of the NC interface.

We conclude that the stabilizing effect of the SEPT9 subunit on filament formation that was already predicted nearly 20 years ago (Sheffield et al. 2003) is mediated by the α6 helix and that this helix is stabilized in its place in the NC interface by the terminal amino acids. Without this stabilizing effect the filament formation properties of octameric rods equal those of hexamers.

### Human septin complexes interfere with actin in the early stages of polymerization

Several studies have already demonstrated the capability of SEPT9, septin hexamers and octamers to bind and bundle filamentous actin (Mavrakis et al. 2014; Iv et al. 2021; Smith, Dolat, Angelis, Forgacs, Spiliotis, and Galkin 2015). All these studies monitored the effects of septins on filamentous actin. Actin bundling was observed subsequently by electron- or fluorescence microscopy. The pyrene actin assay introduced in this study looks at actin polymerization in real time, however it measures the net polymer weight and does not provide information on the longitudinal distribution of the filaments (Cooper, Walker, and Pollard 1983). The assay revealed that the septin rod decreases the rate of polymerization. As we could not detect any interaction between septin rods and G-actin we propose that the septins exert their influence on polymerization through interaction with the forming actin filament.

The following hypothetical model aims at explaining our observations in more detail:

Actin filaments appear as two strands of subunits that wind around each other. The pitch of a single actin strand is 72 nm and consequently 36 nm for the double strand (Milligan, Whittaker, and Safer 1990). The length of eight septin subunits is approximately 35 nm as can be calculated from the available crystal structures of human filamentous septins (PDB-IDs 2QAG and 7M6J), matching almost exactly the pitch of the actin strand. We propose that one octamer aligns per pitch of the growing actin filament. After successful alignment, octamers bundle filaments by lateral joining. This parallel alignment during early stages of polymerization is reflected in our assay by the shift of the inflection point. As the shift is concentration independent for octamers, saturation is likely reached.

The length of one hexamer is about 29 nm. The hexamer does therefore not fit exactly in one pitch of the actin filament. While the sequence of interfaces within an octamer (SEPT2-6-7-9-9-7-6-2) is G-NC-G-NC-G-NC-G it is in the hexamer (SEPT2-6-7-7-6-2) G-NC-G-G-NC-G and thus the order of structural elements that may contribute to actin binding and lateral joining changes considerably. In growing actin filaments, hexamers may align due to their shorter length before a pitch is completed but likely with less and/or more transient affinity and with less bundling capability. This is reflected by the concentration dependency of the hexamer curve shifts in our assay.

Whether SEPT9 is a bundling factor on its own is currently under debate (Mavrakis et al. 2014; Iv et al. 2021; Smith, Dolat, Angelis, Forgacs, Spiliotis, and Galkin 2015). Octamers, but not hexamers have an influence on actin monomer turnover in the elongation phase as detected by our assay. Taken these findings together, we support a contribution of SEPT9 at least in the context of a rod.

### Nucleotide uptake- and hydrolysis by human septin complexes

Septins were first labeled as GTP binding proteins in Drosophila (Field et al. 1996) and afterwards nucleotide binding was confirmed also for human and yeast septin complexes (Sheffield et al. 2003; Farkasovsky et al. 2005). However, the functional role of nucleotide binding- and hydrolysis of the septins in cellular processes and in septin (proto-)filament assembly is still under debate and not well understood.

The availability of yeast and human septin octamers enabled us to measure nucleotide association kinetics in these protein complexes and to compare the properties of both species. GTP-loading in human septin rods remains on a constant level after reaching a plateau whereas in the yeast septin rods the radioactive signal decreases about 50 % after reaching a maximum (Baur et al. 2018). This indicates the release of the γ-phosphate or the entire nucleotide in at least some subunits of the protofilaments. Mg^2+^ suppresses this release but shows no significant effect on the GTP association rate in the yeast septin rods. In contrast, Mg^2+^ slightly slows down the GTP binding in human septin rods. In the absence of Mg^2+^, hexameric and octameric rods from both species are able to exchange previously bound GTP and GDP. The exchange rates for GDP and GTP of the human septin rods do not differ significantly. In contrast, the *S. cerevisiae* septin rods exhibit a much faster GDP-GTP than GTP-GTP exchange. These discrepancies indicate that although displaying a high structural similarity the septins from different species follow each an individual biochemistry. Even within one species the different subunits exhibit different properties. Comparing the GTP and GDP loaded structures of SEPT2 (PDB IDs 3FTQ and 2QNR, respectively) reveals a conformational change in the Switch1 region with the invariant Thr78 and Mg^2+^ being in close contact with the γ-phosphate exclusively in the GTP conformation. In the GDP conformation Switch1 is either unstructured or oriented away from the nucleotide (Suppl. Fig. 5A).

However, this conformational change cannot be seen in the structures of SEPT9 (PDB IDs 5CYP and 5CYO) where the corresponding Thr64 in Switch1 is associated with Mg^+^ towards the nucleotide in in both nucleotide states (Suppl. Fig. 5B). This inflexibility of Switch1 might explain the inability of the isolated SEPT9 to be loaded with [γ^32^P]-GTP and why we do not detect significant differences in nucleotide uptake between hexamers and octamers.

GTP hydrolysis is routinely detected in small GTPases of the Ras family. Experimentally, the small GTPase is loaded with GTP and then subjected to a Mg^2+^containing buffer. The Mg^2+^ ion is an indispensable component for the GTPase reaction. The γ-phosphate is coordinated by invariant Thr and Gly residues in the Switch1 and Switch2 regions. The conformational change following the GTPase reaction was termed “loaded-spring mechanism” which allows the release of the γ-phosphate after hydrolysis and the relaxation of the switch regions into the GDP conformation (Vetter and Wittinghofer 2001).

We were not able to detect the release of [γ^32^P]-GTP or hydrolyzed [γ^32^P] from any rod species examined under these conditions. However, GTPase hydrolysis products were detected in yeast and mammalian septin preparations after denaturation of the protein (Castro et al. 2020; Sirajuddin et al. 2009; Sheffield et al. 2003; Versele and Thorner 2004). In contrast, our assay was conducted under native conditions. This indicates that bound nucleotides and presumable hydrolysis products remain in the binding pocket and are not released into the surrounding medium.

We propose that GTP binding and hydrolysis in septins do not follow the loaded spring mechanism of small GTPases and that septin rods from budding yeast differ mechanistically in their guanine nucleotide interactions from their human counterparts. More structural information from yeast septins is needed to understand these differences.

## 6 Author Contributions

MF, DF and TG performed the experiments. RR performed the MS analysis. NJ analyzed the data and improved the manuscript. TG analyzed the data, conceived the study and wrote the manuscript.

## 7 Conflict of interest

The authors declare no conflict of interest.

## 8 Acknowledgments

The authors thank Matthias Hecht (Inst. f. Molecular Genetics, Ulm) for providing the SEPT9 G domain expression construct and 1306 fibroblast cDNA and Elias Spiliotis (Drexel University, PA, USA) for the kind gift of a GFP-SEPT9i1 expression plasmid. Sebastian Iben and Tamara Pham (Uniklinik Ulm) are acknowledged for providing access and support to the gradient mixing device and Benjamin Grupp (Inst. f. Molecular Genetics, Ulm) for critically reading the manuscript.

## 10 Supplementary Material

Supplementary methods and supplementary results are available as separate files.

## Notes

### Competing Interest Statement

The authors have declared no competing interest.

