## Supplemental Tables and Figures for "Biochemical characterization of a human septin octamer"

**Supporting Information: Supplementary Tables and Figures.**

**Supplementary Table 1**

Evaluation of the actin polymerization reaction. All measurements were performed in triplicate. Y0 is the start value at t=0, $Y_M$ is the maximum value, the rate constant K equals the velocity of actin polymerization, $\tau$ describes the mean doubling time of polymerizing actin and the doubling time $t_{1/2}$ represents the time span after which the initial fluorescence has doubled its value. $t_{ip}$ represents the time point of the inflection point and can be used to compare the time shifts of the individual curves on the x axis.

| | 2 µM actin | + 0.5 µM hexamer | + 1 µM hexamer | + 0.5 µM octamer | + 1µM octamer |
|---|---|---|---|---|---|
| Y0 | 5.77 | 7.32 | 8.46 | 8.38 | 8.65 |
| $Y_M$ | 96.77 | 98.28 | 98.96 | 97.19 | 94.50 |
| K [$s^{-1}$] | 0.001758 | 0.001291 | 0.001262 | 0.001264 | 0.000833 |
| $\tau$ [s] | 569 | 775 | 792 | 791 | 1200 |
| $t_{1/2}$ [s] | 394.4 | 536.9 | 549.2 | 548.5 | 832.1 |
| $t_{ip}$ [s] | 1137.1 | 1518.0 | 2547.7 | 1816.8 | 1861.6 |
| $R^2$ (fit) | 0.9368 | 0.9703 | 0.9476 | 0.9508 | 0.8753 |

**Supplementary Table 2**

Summary of the performed fits for the nucleotide uptake measurements as shown in the figures. $t_{1/2}$ values were calculated as mean from the respective one phase exponential fit. All measurements were performed in triplicate.

| Septin construct | Preload nucleotide | K ($min^{-1}$) | $t_{1/2}$ (min) | 95 % CI of $t_{1/2}$ | $R^2$ (fit) |
|---|---|---|---|---|---|
| Octamer | - | 0,04225 | 16,41 | 15,18 to 17,75 | 0,9974 |
| Hexamer | - | 0,06916 | 10,02 | 7,850 to 12,77 | 0,9759 |
| Octamer + $MgCl_2$ | - | 0,03293 | 21.05 | 18,04 to 24,70 | 0,9916 |
| Hexamer + $MgCl_2$ | - | 0,04794 | 14,46 | 12,32 to 17,02 | 0,9898 |
| Octamer | GDP | 0,02076 | 33,39 | 25,96 to 45,06 | 0,9854 |
| Octamer | GTP | 0,01823 | 38,01 | 32,56 to 45,25 | 0,9953 |
| Octamer | $^{32}$P-GTP | 0,0149 | 46,58 | 40,54 to 54,20 | 0,9957 |
| Hexamer | GDP | 0,01604 | 43,21 | 37,37 to 50,82 | 0,9964 |
| Hexamer | GTP | 0,01684 | 41,15 | 29,73 to 62,65 | 0,9776 |

**Supplementary Table 3**

Evaluation of the nucleotide uptake data sets (each performed in triplicate) with unpaired t-tests. $t_{1/2}$ values were individually calculated for each replicate and provided as triplicate to the statistics analysis.
Pairwise comparisons of the GDP and GTP preloading experiments are only provided for completeness. ANOVA analysis is more applicable for these data.

| | Mean $t_{1/2}$ octamer | Mean $t_{1/2}$ hexamer | Significance | p-value |
|---|---|---|---|---|
| untreated | 16,37 | 10,05 | * | 0,04 |
| +$MgCl_2$ | 21,31 | 14,66 | ** | 0,01 |
| GDP preload | 37,79 | 45,67 | ns | 0,08 |
| GTP preload | 39,39 | 45,96 | ns | 0,40 |
| | Mean $t_{1/2}$ untreated | Mean $t_{1/2}$ + $MgCl_2$ | Significance | p-value |
| Hexamer | 10,05 | 14,66 | ns | 0,13 |
| Octamer | 16,37 | 21,31 | ** | 0,01 |
| | Mean $t_{1/2}$ untreated (min) | Mean $t_{1/2}$ GDP preload (min) | Significance | p-value |
| Hexamer | 10,05 | 45,67 | *** | <0,001 |
| Octamer | 16,37 | 37,79 | ** | 0,003 |
| | Mean $t_{1/2}$ untreated | Mean $t_{1/2}$ GTP preload | Significance | p-value |
| Hexamer | 10,05 | 45,96 | ** | 0,007 |
| Octamer | 16,37 | 39,39 | *** | <0,001 |

**Supplementary Table 4**

Evaluation of nucleotide uptake data for GDP and GTP preloaded constructs (each performed in triplicate) with ordinary ANOVA analysis followed by Holm-Sidak's multiple comparisons test. $t_{1/2}$ values were individually calculated for each replicate and provided as triplicate to the statistics analysis.

| Hexamer | Mean $t_{1/2}$ untreated | Mean $t_{1/2}$ preload | Significance | p-value |
|---|---|---|---|---|
| untreated vs. GDP preload | 10,05 | 45,67 | ** | 0,003 |
| untreated vs. GTP preload | 10,05 | 45,96 | ** | 0,003 |
| | Mean $t_{1/2}$ GDP | Mean $t_{1/2}$ GTP | Significance | p-value |
| GDP vs. GTP preload | 45,67 | 45,96 | ns | 0,96 |
| Octamer | Mean $t_{1/2}$ untreated | Mean $t_{1/2}$ preload | Significance | p-value |
| untreated vs. GDP preload | 16,37 | 37,79 | *** | <0,001 |
| untreated vs. GTP preload | 16,37 | 39,39 | *** | <0,001 |
| untreated vs. $^{32}$P-GTP preload | 16,37 | 46,83 | *** | <0,001 |
| | Mean $t_{1/2}$ GDP | Mean $t_{1/2}$ GTP | Significance | p-value |
| GDP vs. GTP preload | 37,79 | 39,39 | ns | 0,61 |
| | Mean $t_{1/2}$ GTP | Mean $t_{1/2}$ $^{32}$P-GTP | Significance | p-value |
| GTP vs. $^{32}$P-GTP preload | 39,39 | 46,83 | ns | 0,08 |

**Supplementary Figure 1**

**A.** Chromatogram of a SEC purification of octamers containing SEPT9$_{G568}$.The shoulder containing the purified complex is marked with an arrow.
**B.** SDS-PAGE (Coomassie stain) of the marker proteins thyroglobulin (670 kD), γ-globulin (155 kD; separating in SDS-PAGE into 50 kD and 25 kD bands), ovalbumin (44 kD) and myoglobin (17 kD) separated by an 15%-30% Glycerol gradient. Fractions are numbered; B = bottom fraction
**C.** SDS-PAGE (Coomassie staining) of a representative purification of a septin hexamer. IMAC and SEC fractions are indicated.

A

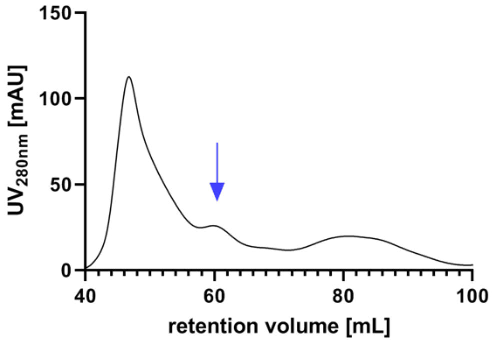

B

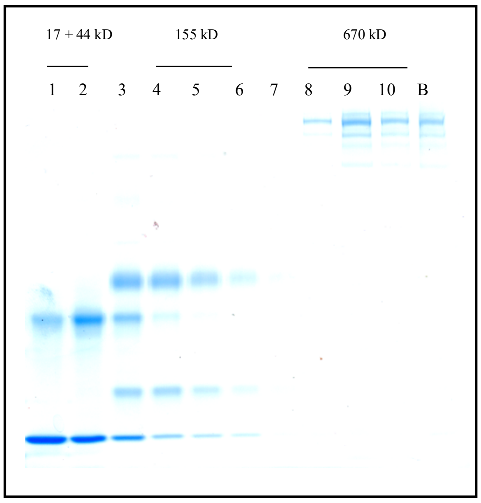

C

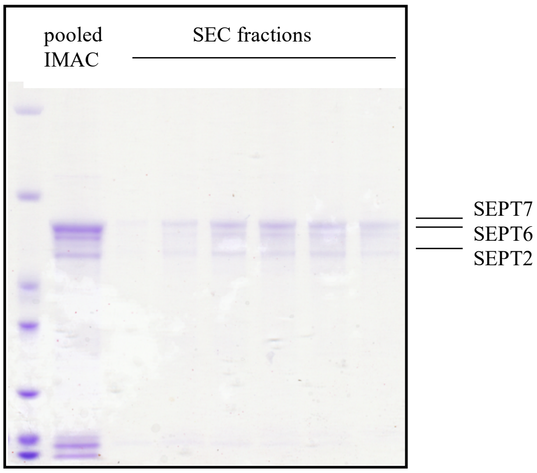

**Supplementary Figure 2**

**A**. Polymerization curve of pyren actin in G buffer. The assay was performed in triplicate.
**B**. Spin down assay of actin (A) and septin octamer (S) in G buffer. Supernatant and pellet were separated on a SDS-PAGE and analyzed by Coomassie-staining. Actin was additionally detected by Western blot using a HRP coupled anti-actin antibody (lower panel). A small fraction of actin aggregates pellets under these conditions.

A

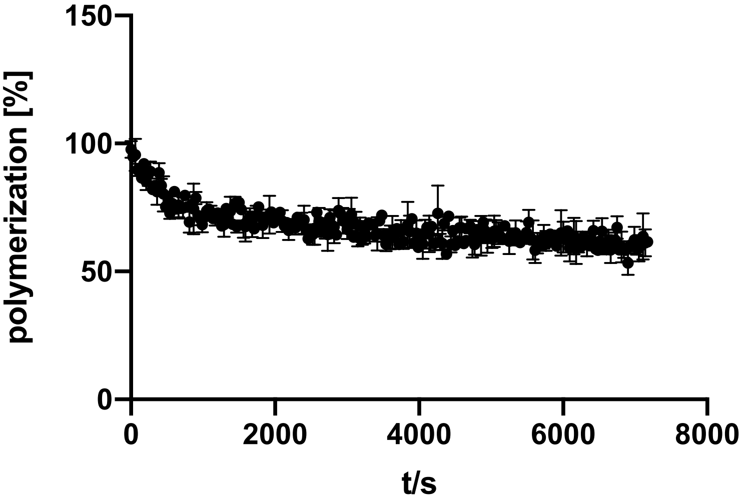

B

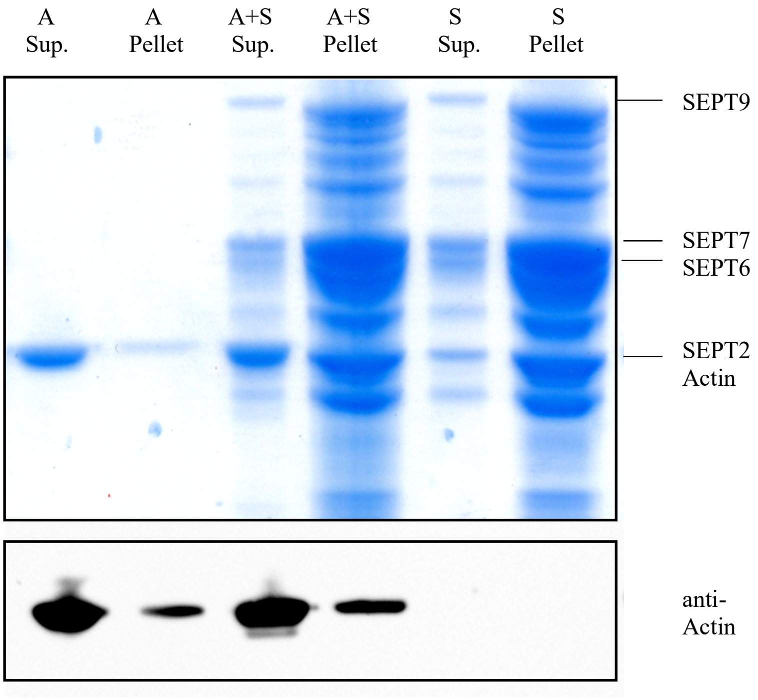

**Supplementary Figure 3**

Nucleotide exchange assays with various conditions. The purified septin complexes were incubated with [$\gamma^{32}$P]-GTP and uptake was monitored at the indicated timepoints by a filter assay. [CPM%] values (normalized radioactive counts) are plotted vs. the reaction time. The connecting line represents the fitting curve of the exponential association. All assays were performed at least in tripliate.

**A.** Nucleotide exchange reaction of hexameric (left panel) and octameric septins (right panel) with $MgCl_2$ added after 20 min.

**B.** Nucleotide exchange reaction of hexameric (left panel) and octameric septins (right panel) preloaded with GDP.

**C.** Nucleotide exchange reaction of hexameric (left panel) and octameric septins (right panel) preloaded with GTP.

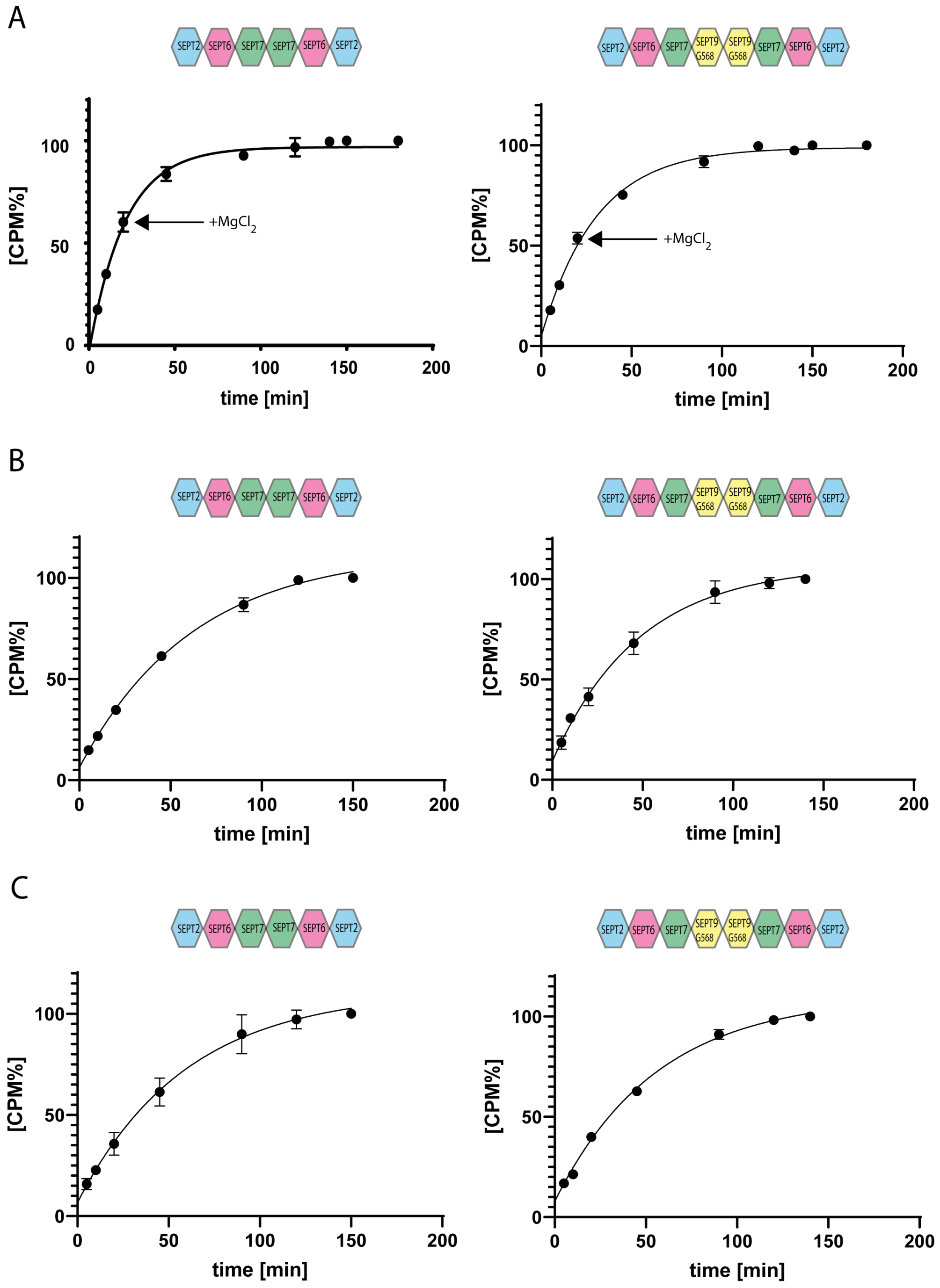

Supplementary Figure 4

Representative GTP hydrolysis assay for hexameric septins. After preloading with [$\gamma^{32}$P]-GTP, the complex was subjected to hydrolysis conditions applicable for small GTPases. The reaction was performed at 25°C.

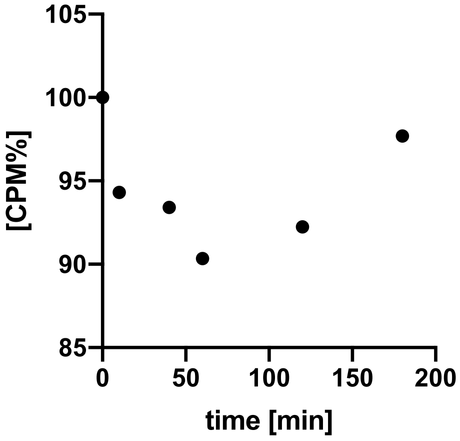

**Supplementary Figure 5**

**A.** Orientation of the Switch1 in GDP (cyan) and GTP (green) bound SEPT2. The nucleotides and Thr64 of Switch1 are shown in stick projection; the black arrow points towards Thr64. The $Mg^{2+}$ ion is shown as green sphere. The figure was derived from PDB-Ids 2QNR and 3FTQ which show both a G-interface SEPT2 dimer. Note that in each structure one SEPT2 subunit has a partially disordered Switch1 and thus the two situations with an ordered Switch1 are shown separately.
**B.** Orientation of the Switch1 in GDP (cyan) and GTP (green) bound SEPT9. The nucleotides and Thr64 of Switch1 are shown in stick projection. The $Mg^{2+}$ ion is shown as green sphere. Note the well superimposed Switch1 in both structures. The figure was derived from PDB-IDs 5CYO and 5CYP which show both a G-interface SEPT9 dimer.

A

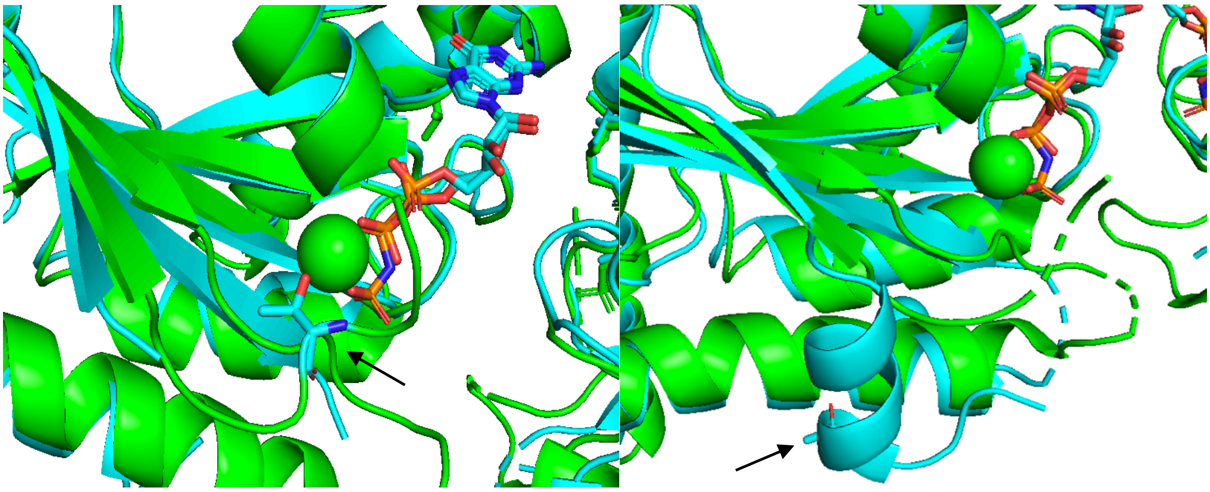

B

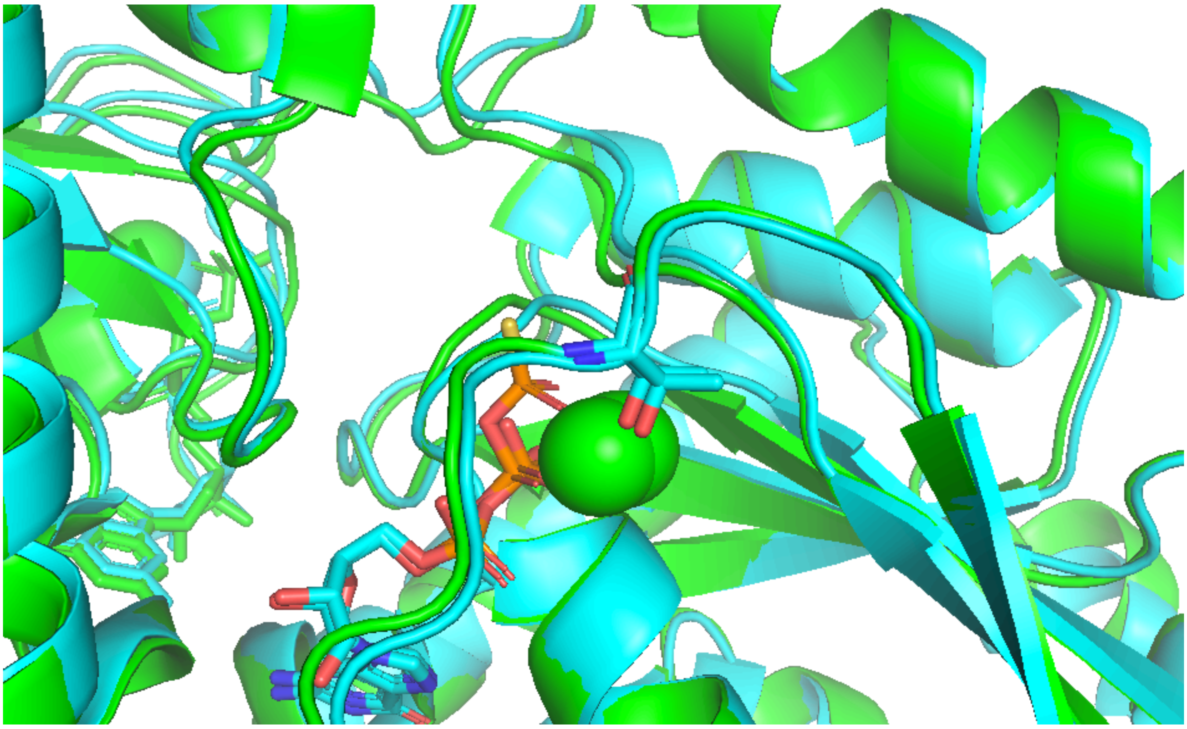
