## Supplemental Methods for "Biochemical characterization of a human septin octamer"

**Supporting information: Supplementary methods**

*Cloning of the expression constructs*

All generated plasmids are summarized in Supplementary Methods Table 1.

The ORFs of the isoforms i1 of SEPT2 and SEPT9 (or SEPT9$_{1-568}$) were cloned into the bicistronic plasmid pACYC-Duet (Novagen). SEPT7 and SEPT6 were cloned into the compatible plasmid pET-Duet1 (Novagen), the latter in frame downstream of the 6-his tag. SEPT2 and SEPT6 were PCR amplified from cDNA obtained from human 1306 fibroblast cells. SEPT9 was PCR amplified from a GFP-SEPT9i1 expression plasmid (kind gift of E. Spiliotis, Drexel University, PA, USA) and plasmid #4 containing SEPT7 was constructed via gene synthesis. For the expression of hexameric rods we used SEPT2 in pACYC (plasmid #1) without inserted SEPT9.

6his-SEPT9 and the 6his-SEPT9 G domain were obtained by cloning the respective PCR amplificates into an in-house constructed pET15b derivative allowing for the expression of N-terminally tagged fusion proteins.

Used primers are listed in Supplementary Table 2.

Supplementary Methods Table 1 – Generated expression constructs. Cloning plasmid pES is an in-house constructed pET15b derivative.

| | Insert | Cloning plasmid | Restriction sites | Ori | Combination plasmid for expression |
|---|---|---|---|---|---|
| #1 | SEPT2 | pACYC-Duet | Eco1, Sal1 | p15A | |
| #2 | SEPT2+SEPT9 | #1 | Mfe1, Xho1 | p15A | #5 for octamer |
| #3 | SEPT2+SEPT9$_{M1-G568}$ | #1 | Mfe1, Xho1 | p15A | #5 for SEPT9$_{G568}$ octamer |
| #4 | SEPT7 | pET-Duet1 | Mfe1, Xho1 | pBr322 | |
| #5 | His-SEPT6+SEPT7 | #4 | BamH1, Hind3 | pBr322 | #1 for hexamer |
| #6 | His-SEPT9 | pES | Sfi1 | pBR322 | |
| #7 | His-SEPT9$_{Q295-E567}$ | pES | Sfi1 | pBR322 | |

Supplementary Methods Table 2 – Primer sequences

| Name | Sequence 5' -> 3' (restriction sites are underlined) |
|---|---|
| Sept2-EcoR1for | gcatgaattcatgtctaagcaacagccaactc |
| Sept2stop-Sal1rv | gcgtcgactacacgtggtgcccgagag |
| Sept9-Mfe1for | gctgacaattggatgaagaagtcttactcaggag |
| Sept9stop-Xho1rv | cgatctcgagctacatctccggggcttctg |
| Sept9-G568stop-Xho1rv | cagtctcgagctagccctcgttgaggcgcttc |
| Sept9(Q295)-Sfi1for | gcatcggccagcacggcccagggcttcgagttcaacatcatg |
| Sept9-E567stop-Sfi1rv | catgaggccaaaaaggcctcactcgttgaggcgcttcacac |
| Sept9-Sfi1for | ccgaaggccagcacggccgaaaacctgtacttccagggtaaaaagtcttactcaggagg |
| Sept9-Sfi1rv | ctgtgggccaaaaaggccttatcacatctctggggcttctgg |
| Sept6-BamH1for | catgggatccgatggcagcgaccgatatagc |
| Sept6stop-Hind3rv | gatcaagcttttaattcttcttctctttgtctctc |

*Protein expression and purification*

Protein expression of hexameric and octameric rods was performed in the *E. coli* strain BL21DE3. The combination of plasmids for these protein complexes are outlined in Supplementary Methods Table 1. Cells were grown in SB medium until the $O_D$(600nm) reached 1.0. Protein expression was induced with 1 mM IPTG and the culture was adjusted to 2 % Ethanol. Protein expression was subsequently carried out over night at 18 °C. The pelleted cells were resuspended in buffer IMAC-A (50 mM Tris-HCl pH 8.0, 500 mM NaCl, 2 mM $MgCl_2$, 15 mM imidazole, 5 mM β-Mercaptoethanol, 12 % (v/v) glycerol), supplemented with 0.1 % (v/v) Tween20 and Complete Protease Inhibitor Cocktail (Roche). Cell lysis was performed by treatment with lysozyme (1.0 mg/ml, 30 min on ice) and sonication (Branson Sonifier A250). The resulting lysate was clarified by centrifugation at 40,000 g for 10 min at 4 °C and the supernatant was loaded onto a 5 ml HisTrap HP column (GE Healthcare) mounted on an Äkta Purifier Chromatography System (GE Healthcare). The proteins were eluted from the column by applying consecutive Imidazole segment gradients with 3 %, 10 % and 100 % buffer IMAC-B (50 mM Tris-HCl pH 8.0, 500 mM NaCl, 2 mM $MgCl_2$, 250 mM Imidazole, 5 mM β-Mercaptoethanol, 12 % (v/v) glycerol). The septin rods eluted at 100 % IMAC-B. Septin rod-containing fractions were pooled and subjected to size exclusion chromatography (SEC) against exchange buffer (25 mM Tris-HCl pH 8.0, 300 mM NaCl) or high salt buffer (25 mM Tris-HCl pH 8.0, 500 mM NaCl) using a Superdex 200 16/60 column (GE Healthcare). Protein concentrations were determined at 280 nm in a NanoDrop ND-1000 spectral photometer (Peqlab) with calculated extinction coefficients or with a Bradford assay. Bioinformatic tools provided on the EXPASY server were used for calculating molecular weights and extinction coefficients.

*Density gradient centrifugation*

Integrity of the isolated octameric complex was assayed by applying the IMAC eluate on a 15% - 30% Glycerol gradient in 25 mM Tris pH 8.0, 300 mM NaCl which was prepared using a Gradient Station IP gradient mixing device (Biocomp Instruments). The gradient was separated by ultra-centrifugation in a Optimax-XP centrifuge (Beckman) in a MLS-50 swing out rotor (Beckman) at 268.000 xg for 10:30h. The gradient was separated into 11 fractions and assayed by SDS-PAGE and Coomassie staining. A gel filtration standard protein mix (Biorad) composed of thyroglobulin (670 kD), γ-globulin (155 kD; separating in SDS-PAGE into 50 kD and 25 kD bands), ovalbumin (44 kD) and myoglobin (17 kD) was used as size standard.

*Mass spectrometry*

4 µg of total protein (i.e. septin octamer from a SEC purification) were reduced with 5 mM DTT for 20 min at RT, alkylated with iodoacetamide for 20 min at 37 °C and diluted with 50 mM ammonium bicarbonate. Trypsin was added in a 1:50 enzyme-protein ratio and digested overnight at 37 °C. Peptides were cleaned up using self-made C18-STAGE tips (Empore C18 Extraction Disk, 3M) and following evaporation dissolved in 15 µl 0.1% TFA.

Samples were measured using an LTQ Orbitrap Elite system (Thermo Fisher Scientific) online coupled to an U3000 RSLCnano (Thermo Fisher Scientific) as described previously (Hecht et al. 2019) with the following modifications: After loading and washing, the peptides were transferred to the analytical column and eluted by raising the percentage of B from 5 to 15% in 5 min, followed by an increase of 15 to 40% B in 145 min. The eluted peptides were ionized and introduced into an LTQ Orbitrap Elite mass spectrometer using a nanospray source. Full scans ranging from m/z 370 to 1700 were acquired in the Orbitrap at a resolution of 30,000 (at m/z 400) with Automatic gain control (AGC) enabled. The 20 most intense ions from the survey scan were picked for CID fragmentation, with collision energy set to 35% and an activation Q

of 0.25. Singly charged ions were rejected and m/z of fragmented ions were excluded from fragmentation for 60 s. MS2 spectra were acquired employing the LIT at rapid scan speeds.
Database search was performed using MaxQuant Ver. 1.6.3.4 (www.maxquant.org) (Cox and Mann 2008). For peptide identification the build-in Andromeda search engine (Cox et al. 2011) was employed to correlate MS/MS spectra with the respective septine protein sequences obtained www.uniprot.org). Carbamidomethylated cysteine was considered as a fixed modification along with oxidation (M), and acetylated protein N-termini as variable modifications. False Discovery rates were set on both, peptide and protein level, to 0.01.
Protein intensities were calculated as the sum of all identified peptide intensities (unique and razor peptides) belonging to the respective protein group.

*Actin polymerization assays*

Pyrene-labeled G-actin (shortly pyrene actin) was prepared from rabbit muscle acetone powder using an already described protocol (Cooper, Walker, and Pollard 1983) with some modifications: 3.5 g acetone powder were resuspended in 125 ml ice cold G buffer (25 mM Tris pH 8.0, 0.2 mM ATP, 0.5 mM DTT, 0.1 mM $CaCl_2$) and stirred for 2 h at 4 °C to extract actin. Cell debris was separated by centrifugation and the supernatant was filtered through a Whatman filter, adjusted to 15 mM NaCl and 2 mM $MgCl_2$ and stirred overnight at 4 °C. The next day NaCl was adjusted to final 0.6 mM by adding NaCl powder and actin polymerization was allowed to continue for further 30 minutes.
F actin was collected by centrifugation (2.5 h, 100,000 g) and allowed to depolymerize again by dounce homogenization and subsequent dialysis in G buffer for 48 hrs.
G-actin was cleared from polymerized remnants by centrifugation and dialyzed against G Buffer without DTT for 4 h. G-actin as adjusted to 23.5 mM in labeling buffer (25 mM Tris pH 7.5, 100 mM KCl, 0.3 mM ATP, 2 mM $MgCl_2$) and 7-fold molar excess N-(1-Pyrenyl)-iodoacetamide (ChemImpex Int.) (dissolved in DMF) was added dropwise and the solution was rotated overnight at 4 °C protected from light. Precipitated pyrene dye was removed by centrifugation at 4000 rpm in a benchtop centrifuge and labelled F-actin was collected by centrifugation at 100,000 x g for 3 h. The pellet was collected in G buffer, dounced and dialyzed into G buffer for 48 h at 4 °C.
G-actin was cleared from polymerized remnants by centrifugation and subjected to size exclusion chromatography on a Superdex 200 16/60 column (GE Healthcare) against G buffer. Peak fractions with a labeling efficiency (determined by the $OD_{344}$/ $OD_{290}$ ratio) of greater than 5 % were pooled, concentrated and used for the polymerization assays.
Septin octamer or hexamer were prepared in suitable concentrations in 10 µl 10x polymerization buffer (25 mM Tris pH8, 300 mM NaCl, 10mM $MgCl_2$). The reaction was started by adding 90 µl 2.2 µM pyrene actin, freshy diluted in water, and the polymerization kinetics were recorded in a Tecan Spark plate reader for 2 h with the excitation set to 365 nm and emission set to 407 nm. Control measurements were performed without septin either in polymerization buffer or in G buffer. The obtained raw response unit values were converted to relative values by scaling to the maximum value which was set to 100%.
Actin polymerization is characterized by a lag period ($y_0$) where G-actin forms nuclei setting off the reaction and entering an exponential phase of filament growth. As G-actin monomers are incorporated into the growing filament and free G-actin concentration decreases, the curve levels off and reaches an asymptotic plateau phase. The resulting plot resembles a sigmoidal curve which was fitted with the nonlinear regression model "sigmoidal 4PL," in Prism (GraphPad Software), following equation (1).

$$(1)\ y(t) = \frac{Top - Bottom}{1+\left(\frac{IC50}{t}\right)^{h}} + Bottom$$

h is the hill slope (representing the steepness of the graph), IC50 is the point of time at which intensity reaches half-maximum value, Top is the maximum fluorescence intensity and Bottom the minimum fluorescence intensity, respectively. The inversion point of the sigmoidal graph is calculated by the second derivative equaling zero (equation (2)). Inserting the x-value of the inversion point $t_{ip}$ into the first derivative y'($t_{ip}$) yields the slope at the inversion point. The calculated properties were substituted in equation (1), then derivatized two times with Wolfram Mathematica (https://www.wolfram.com/mathematica/) to calculate the inversion point of the sigmoidal graph.

$$(2)\ y''(t) = 0 \text{ for } t = t_{ip}$$

The inversion point provides insight into the time shift of curves relative to each other, thus providing information on the length of the lag period.
Subsequently, all values on the time axis prior to the point of inversion were deleted and the nonlinear regression model "One Phase Association" (equation (3)) was applied to obtain the rate constant K (in $s^{-1}$), doubling time $t_{1/2}$ (in s) and the time constant $\tau$ (in s, equation (6)) of the elongation reaction. The doubling time is calculated as ln(2)/k and the time constant represents 1/K. Y0 is the start value at t=0 and $P_{lateau}$ is the Y value at infinite times.

$$(3)\ y(t) = (Plateau - y(t_{ip})) \cdot (1 - e^{Kt}) + y(t_{ip})$$

X-fold increase or decrease of parameters compared to the respective actin blank measurement dependent on each pyrene actin purification approach was computed.

To determine if septin filaments interact with G-actin, septin octamer was dialyzed into G buffer without DTT and ATP. Unlabeled G actin was cleared from aggregates by centrifugation for 20 min at 280,000 g, set to 2 µM and incubated with 2 µM septin octamer in G or F buffer for 105 min. After centrifugation for 20 min at 280,000 g, pellet and supernatant were analyzed by SDS-PAGE followed by Coomassie staining or Western blot using an anti-actin HRP conjugate (Santa Cruz Biotechnology).

*Nucleotide uptake- and hydrolysis assays*

Uptake of [$\gamma^{32}$P]-GTP to hexameric and octameric rods and isolated SEPT9 was performed as described elsewhere for yeast septins (Baur et al. 2018). Briefly, septin preparations were incubated with [$\gamma^{32}$P]-GTP in exchange buffer containing 5 mM EDTA and 1 mM DTT. For hydrolysis, the exchange reaction was transferred into hydrolysis buffer containing 5 mM $MgCl_2$ and 10 µM non-radioactive GTP. Radioactivity retained in the native septin proteins during uptake or during hydrolysis was detected with a filter binding assay.
Dynamics of the nucleotide exchange was monitored by loading hexameric or octameric $SEPT9_{G568}$ containing rods with GDP, GTP or [$\gamma^{32}$P]-GTP for 45-60 min. Excess nucleotide was removed by micro-dialysis on a 0.025 µM membrane (VSWP nitrocellulose, Merck-Millipore) or by passing the mixture over a NAP5 desalting column (GE Healthcare), respectively. The preparation was subsequently subjected again to an exchange reaction with the desired nucleotide as described above. All assays were performed in triplicate.
The raw CPM counts were prior to evaluation converted to relative CPM% values by scaling them to the maximum value which was set to 100% (Baur et al. 2018).
[$\gamma^{32}$P]-GTP uptake followed an exponential one phase association kinetics which can mathematically be described by aquation (4).

$$(4)\ y(t) = Y0 + (Plateau - Y0) \cdot (1 - e^{Kx})$$

Dissociation kinetics can be described by an exponential one-phase decay following equation (5).

$$(5)\ y(t) = (Y0 - Plateau) \cdot e^{-Kx} + Plateau$$

Y0 is the start value at t=0, K is the rate constant of the reaction and $P_{lateau}$ is the Y value at infinite times. The reaction half time $t_{1/2}$ is computed as ln2/K for both kinetics. Where individual $t_{1/2}$ values were compared, they were statistically evaluated for significance with an unpaired t-test. If values within a whole group were compared (i.e. all $t_{1/2}$ values of nucleotide uptake for octamers), an ordinary ANOVA test was performed. Significances were allocated according to NEJM standard.
